# Characterization of *Rosa damascena* Callus-derived Exosome-like Vesicles and Their Multifunctional Activities in Skin-related Cellular Models

**DOI:** 10.1101/2023.10.17.562840

**Authors:** Byong Seung Cho, Hyun Ju Lee, Bogeun Son, Esther Lee, Sang Yun Moon, Ella Shin, Jeong Jin Lee, Jun Young Hur, Seong Kyu Park, Cholhyun Park, Kyung-Min Lee, Dae Hyun Ha, Mun Seog Chang

**Affiliations:** ExoCoBio Exosome Institute (EEI), ExoCoBio Inc., Seoul 08594, Republic of Korea; (B.S.C.); (H.J.L.); (B.S.); (E.L.); (S.Y.M.); (E.S.); (D.H.H.); Department of Prescriptionology, College of Korean Medicine, Kyung Hee University, Seoul 02447, Republic of Korea; (J.J.L.); (J.Y.H.); (S.K.P.); (C.P.); (M.S.C.); Department of Life Science, Hanyang University, Seoul 04763, Republic of Korea; Natural Science Institute, Hanyang University, Seoul 04763, Republic of Korea; Hanyang Institute of Bioscience and Biotechnology, Hanyang University, Seoul 04763, Republic of Korea; (K.L.)

**Author notes:** Correspondence (M.S.C.); (D.H.H.).

**Keywords:** *Rosa damascena*, Callus-derived exosome-like vesicles, Plant-derived extracellular vesicles (PDEVs), Skin regeneration, Anti-inflammatory activity, Melanogenesis, Proteomics

## Abstract

Plant-derived extracellular vesicles (PDEVs) are emerging as promising bioactive materials for biomedical and dermatological applications. In this study, we isolated and characterized exosome-like vesicles derived from *Rosa damascena* callus culture medium (RSC-EXO) and evaluated their molecular features and biological activities in skin-related cellular models. Nanoparticle tracking analysis and cryo-electron microscopy showed that RSC-EXO exhibited a nanoscale size distribution and spherical morphology. Western blotting confirmed enrichment of the plant EV-associated markers PEN1 and TET8. RSC-EXO were efficiently internalized by human dermal fibroblasts and showed markedly improved biocompatibility compared with crude conditioned medium (RSC-CM). Functionally, RSC-EXO significantly increased collagen synthesis and showed a trend toward enhanced wound closure in fibroblasts. In addition, RSC-EXO reduced melanin production in α-MSH-stimulated B16F10 melanoma cells and suppressed the secretion of pro-inflammatory cytokines, including IL-1α, IL-6, and TNF-α, in LPS-stimulated RAW 264.7 macrophages. Proteomic analysis revealed a distinct cargo enriched in stress-, defense-, and metabolism-related proteins, while small RNA sequencing identified a heterogeneous small RNA population containing a limited fraction of miRNA-sized reads. Collectively, these findings suggest that RSC-EXO represents a biologically active plant-derived vesicle population with regenerative and anti-inflammatory activity observed *in vitro* in skin-related cellular models and support its potential as a promising platform for future cosmeceutical and dermatological applications.

## 1. Introduction

Exosomes are nano-sized extracellular vesicles (EVs) that mediate intercellular communication by transporting various molecular cargos, such as proteins, lipids, and RNAs [1]. Mammalian exosomes have been extensively studied and applied for diagnostic, therapeutic, and aesthetic purposes owing to their ability to modulate various cellular functions. Among these, adipose-derived stem cell exosomes (ASC-EXOs) have been particularly well characterized and are clinically known to improve skin hydration, barrier function, whitening, and wrinkle reduction [2–8].

Building on these findings, growing attention has recently been directed toward plant-derived extracellular vesicles (PDEVs), which are secreted from various plant tissues including roots, fruits, pollen, and calli [9, 10]. PDEVs offer several advantages over mammalian-derived EVs, including favorable biocompatibility, a generally low immunogenic profile, and the potential for scalable production. They are also being explored as delivery platforms for both oral and transdermal administration [9, 11–13]. In addition, PDEVs have been proposed to mediate inter-kingdom communication, enabling the transfer of bioactive molecules from plants to mammalian cells [14, 15].

Recent studies have demonstrated that PDEVs participate in intercellular signaling and deliver functional molecules such as small RNAs and proteins. They have shown promising roles in diverse physiological and pathological processes, including tissue regeneration, immune modulation, antioxidation, and even antitumor activity. Due to their biocompatibility and ability to carry bioactive molecules, PDEVs have attracted increasing interest for their potential interactions with mammalian systems, particularly as nanocarriers and therapeutic agents [9, 16, 17].

Characteristic plant exosomal markers such as PEN1 and TET8 have been used to define and classify these vesicles. TET8, a plant tetraspanin protein, has been reported as a marker of extracellular vesicles derived from the multivesicular body (MVB)–endosomal pathway and is considered analogous to the mammalian exosomal marker CD63. TET8-positive EVs have been reported to carry small RNAs involved in plant defense responses, including interactions with pathogens [18–20]. PDEVs also contain RNA-binding proteins (e.g., AGO1 and DEAD box helicases), which are thought to regulate selective loading and functional expression of small RNAs [21]. Although plant-derived RNAs have been proposed to regulate gene expression in recipient mammalian cells, their functional relevance in mammalian systems remains controversial and requires further validation and should therefore be interpreted cautiously [22–25].

Given their biocompatibility and regenerative potential, PDEVs are emerging as promising bioactive materials for cosmetic and dermatological applications, with the ability to exert biological effects in mammalian systems and serve as active ingredients in cosmeceuticals or therapeutics [26–28].

Callus represents a dedifferentiated mass of plant cells with stem cell-like properties, and vesicles derived from callus are therefore considered plant stem cell-derived EVs [29, 30]. In particular, plant callus-derived extracts and materials have been reported to exert various dermatological benefits, including antioxidation, moisturization, wound healing, anti-aging, and cell regeneration [31–33]. However, most of these studies have focused on crude extracts or conditioned media rather than purified extracellular vesicles, leaving the molecular basis of callus-derived extracellular vesicle functions largely unexplored, particularly in *Rosa* species.

Building on these findings, *Rosa damascena*-derived ingredients have attracted attention for skin-related applications, and recent studies have reported antioxidant, whitening, and anti-skin-aging effects of *Rosa damascena* extracts in experimental systems [34]. These findings provide supportive context for exploring Rosa-derived materials as potential bioactive ingredients for dermatological and cosmeceutical applications. Moreover, *Rosa*-derived ingredients have been shown to activate Nrf2 and LAMP2A pathways, which are associated with anti-aging and cellular stress response modulation [35].

Recent preclinical and clinical studies have further revealed that Rose stem cell-derived exosomes-like vesicles may promote wound healing and scar reduction, enhance skin regeneration, and even support hair growth, underscoring their translational potential in dermatology [36–39]. However, these studies primarily focused on functional outcomes rather than detailed molecular characterization.

Despite these promising findings, several critical challenges remain, including the heterogeneity of plant-derived vesicles depending on plant species, tissue origin, and isolation methods, as well as the lack of standardized markers and characterization criteria. Unlike conventional single-molecule natural compounds, plant-derived extracellular vesicles represent a complex bioactive system that encapsulates multiple functional components, including proteins, lipids, and RNAs, thereby enabling integrated biological effects. In this context, PDEVs can be considered a natural nano-scale bioactive platform rather than a single isolated compound.

In this study, we aimed to isolate and characterize exosome-like vesicles derived from *Rosa damascena* callus cultures and to investigate their molecular composition and biological activities relevant to human skin. These vesicles are referred to as Rose Stem Cell-derived Exosome-like Vesicles (RSC-EXO) based on the detection of plant exosome-associated markers such as TET8, although the classification and nomenclature of plant extracellular vesicles remain under active discussion. This study provides a comprehensive molecular and functional characterization of RSC-EXO, thereby addressing an important gap in understanding the molecular basis and functional properties of plant-derived extracellular vesicles.

## 2. Results

### 2.1. Isolation and Characterization of RSC-EXO

RSC-EXO were isolated from the callus culture medium of *Rosa damascena* and characterized by nanoparticle tracking analysis (NTA) and cryo-electron microscopy. NTA revealed a mean particle size of 126.9 nm with a mode size of 107.1 nm (Figure 1a). Cryo-electron microscopy confirmed the presence of spherical vesicles ranging from 30 to 200 nm in diameter (Figure 1b).

**Figure 1.**
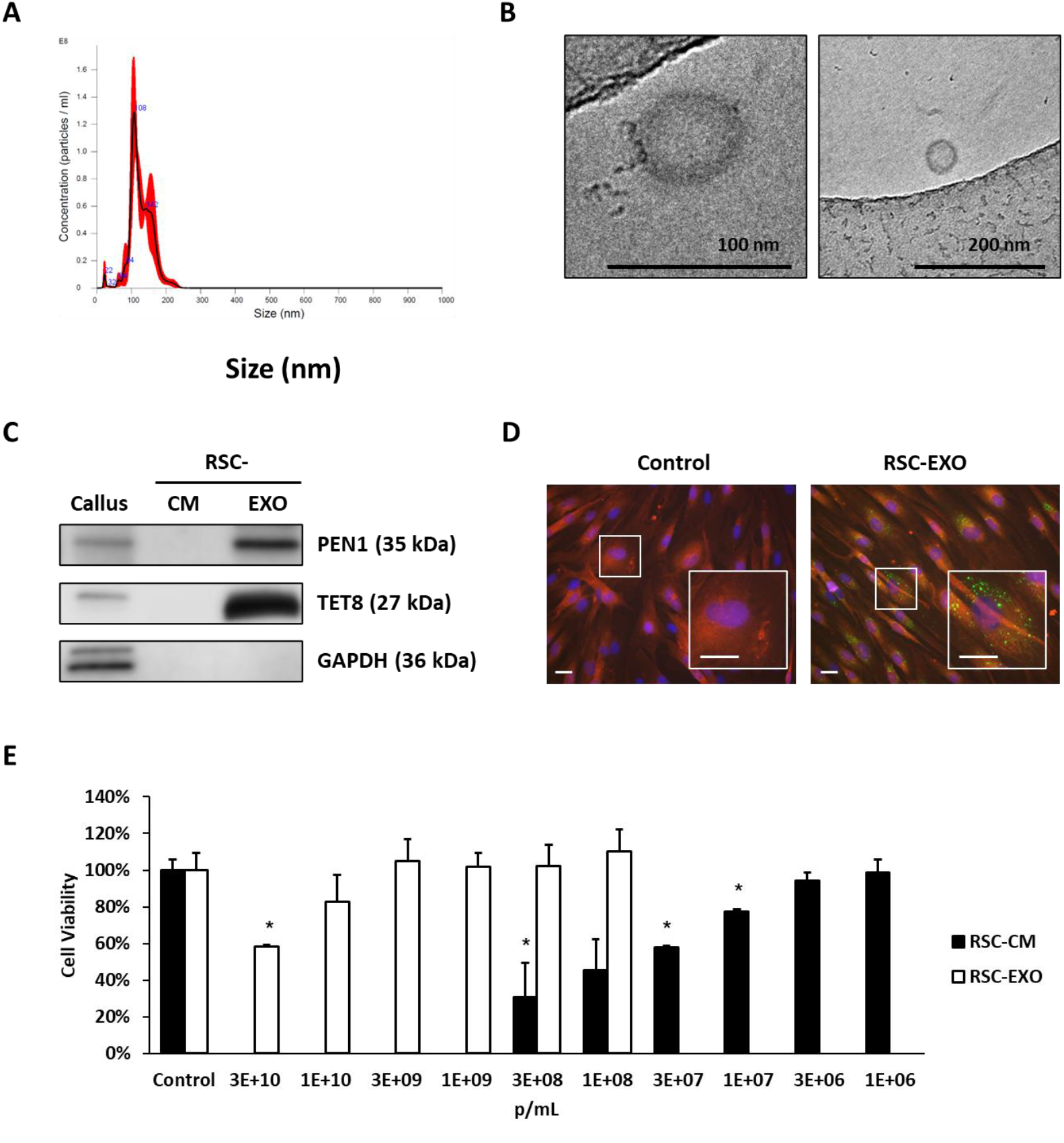
Characterization, cellular uptake, and biocompatibility of RSC-EXO. (A) Representative histogram of the concentration and size distribution of RSC-EXO measured by nanoparticle tracking analysis. (B) Representative cryo-electron microscopy image of RSC-EXO. Scale bars are indicated in the image. (C) Western blot analysis of RSC-EXO for plant exosome-like vesicle markers PEN1 and TET8, and the absence of the cytosolic protein GAPDH. (D) Uptake of PKH67-labeled RSC-EXO by human dermal fibroblasts (HDFs) observed using fluorescence microscopy. Nuclei were counterstained with Hoechst 33258 (blue), and cytoplasm was stained with CellMask (red). A dye-only control (PBS subjected to the same labeling and procedure) showed no detectable fluorescence signal. Scale bar = 10 μm. (E) Cytotoxicity of RSC-EXO and RSC-CM in HDFs. Data are presented as mean ± SD (n = 3). Statistical significance was determined relative to control (\**p* < 0.05, \*\**p* < 0.01).

Western blot analysis demonstrated enrichment of plant EV-associated markers, including PEN1 and TET8, in RSC-EXO compared to conditioned medium (CM), while the cytosolic marker GAPDH was not detected (Figure 1c), suggesting reduced contamination from intracellular components. The plant-associated markers used in this study are based on homologous proteins identified in model plant systems, and their specificity in *Rosa damascena* has not been fully validated.

### 2.2. Cellular Uptake and Biocompatibility of RSC-EXO

To evaluate cellular interaction, uptake of PKH67-labeled RSC-EXO was assessed in human dermal fibroblasts (HDFs). Fluorescence microscopy indicated internalization of RSC-EXO into the cytoplasm (Figure 1d). Importantly, no detectable fluorescence signal was observed in cells treated with the dye-only control processed in parallel, indicating that the observed intracellular fluorescence was not attributable to residual dye or micelle artifacts.

Cytotoxicity was assessed in comparison with RSC-CM. While RSC-CM began to induce measurable cytotoxicity above 1 × 10^7^ particles/mL, RSC-EXO showed no statistically significant cytotoxicity up to 3 × 10^9^ particles/mL, although cytotoxicity became apparent at 3 × 10^10^ particles/mL (Figure 1e).

When normalized by particle number, RSC-EXO exhibited substantially lower cytotoxicity than RSC-CM, supporting its favorable biocompatibility. Notably, achieving an equivalent particle dose required a substantially lower treatment volume for RSC-EXO due to its concentrated form. In contrast, larger volumes of RSC-CM were necessary to deliver the same number of particles. These findings highlight the practical advantage of RSC-EXO as a concentrated vesicle formulation with improved biocompatibility compared with crude conditioned medium.

Based on these findings, a concentration of up to 1 × 10^9^ particles/mL was used for subsequent in vitro experiments.

### 2.3. RSC-EXO Promotes Collagen Synthesis and Wound Closure

Building on the established dermatological benefits of *Rosa damascena*-derived materials, including their roles in antioxidation, anti-inflammatory responses, and skin regeneration, we next sought to evaluate whether RSC-EXO could directly modulate key processes associated with skin repair. In particular, collagen synthesis and wound closure are critical indicators of dermal regeneration and are widely used to assess the functional activity of bioactive skin materials.

To this end, the effects of RSC-EXO on collagen production were examined in human dermal fibroblasts (HDFs). Treatment with RSC-EXO resulted in increased collagen production at both tested concentrations, with 43% and 127% elevation observed at 3 × 10^8^ and 1 × 10^9^ particles/mL, respectively (Figure 2a). In contrast, RSC-CM did not induce comparable effects under the same conditions.

**Figure 2.**
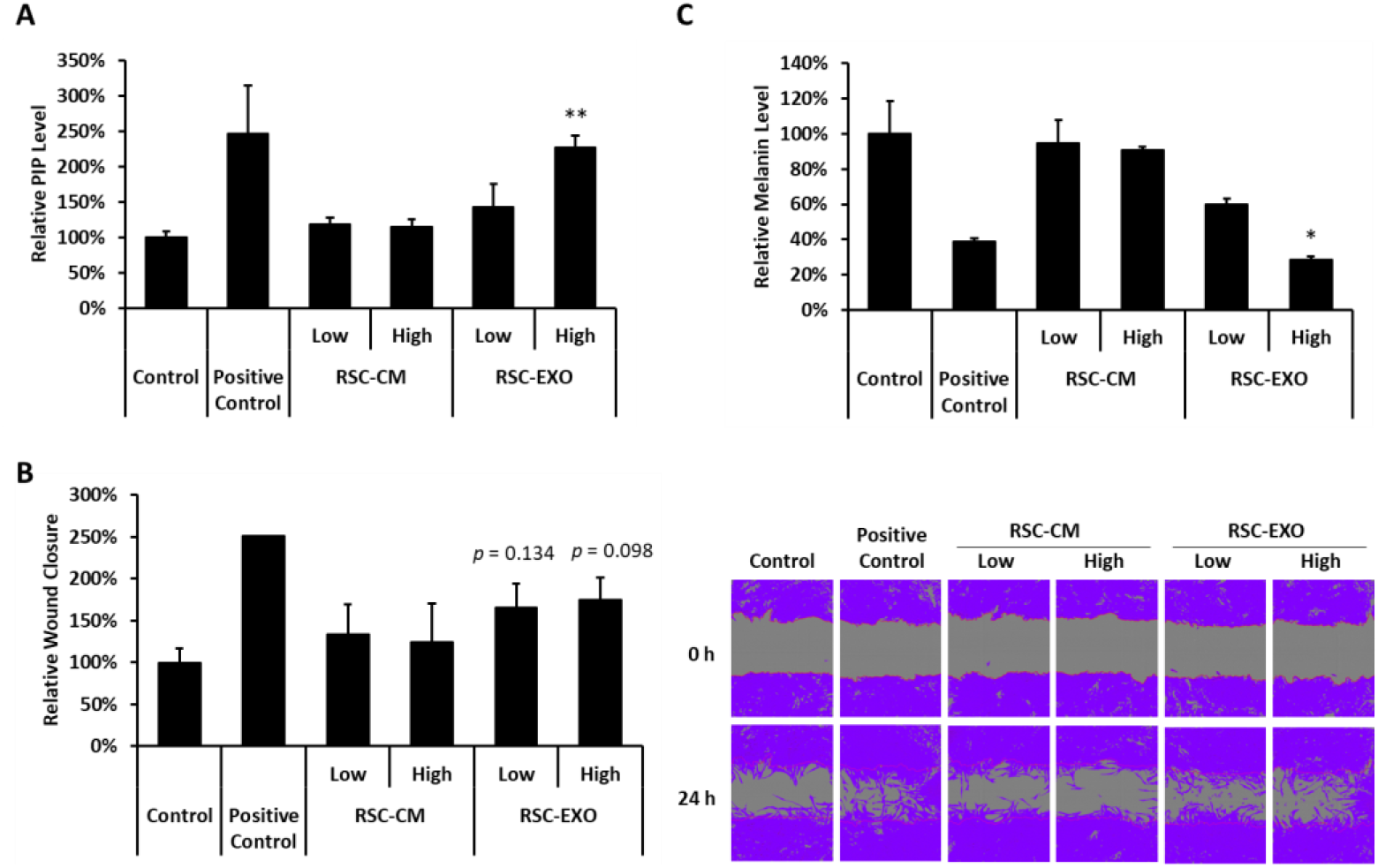
Effects of RSC-EXO on collagen synthesis and wound closure. (A) HDFs were treated with RSC-EXO or RSC-CM, and procollagen type I production was quantified by ELISA. RSC-EXO was applied at high (1×10^9^ particles/mL) and low (3×10^8^ particles/mL) concentrations, while RSC-CM was treated at equivalent volumes. A 5% FBS growth medium was used as a positive control. (B) HDFs were treated with RSC-EXO or RSC-CM, followed by wound closure assays. RSC-EXO was applied at high (1×10^9^ particles/mL) and low (3×10^8^ particles/mL) concentrations, while RSC-CM was treated at equivalent volumes. A 5% FBS medium was used as a positive control. (C) Effects of RSC-EXO on melanin production in α-MSH-stimulated B16F10 melanoma cells. Cells were treated with RSC-EXO or RSC-CM for 48 h in the presence of α-MSH (100 nM). Melanin content was quantified using a standard curve and normalized to cell viability measured by CCK-8 assay. Arbutin (1 mM) was used as a positive control. Data are presented as mean ± SD (n = 3). Statistical significance was indicated as follows: \**p < 0.05*, \*\**p < 0.01*.

The impact of RSC-EXO on cellular migration and wound repair was further assessed using a scratch assay. RSC-EXO treatment resulted in an increase in wound closure at both tested concentrations, reaching 66% and 75% closure at 24 h (Figure 2b), whereas RSC-CM showed no detectable effect. However, these increases did not reach statistical significance (p = 0.134 at 3 × 10^8^ particles/mL and p = 0.098 at 1 × 10^9^ particles/mL), although a consistent trend toward enhanced wound closure was observed.

Taken together, these findings suggest that RSC-EXO enhances collagen synthesis and may contribute to wound closure under the tested conditions, with the latter showing a consistent but not statistically significant trend. These results are consistent with the observed biocompatibility of RSC-EXO and support its potential as a functional skinregenerative material.

### 2.4. RSC-EXO Modulates Melanin Production in Melanoma Cells

In addition to dermal regeneration, modulation of pigmentation represents another important aspect of skin physiology and cosmetic relevance. Therefore, we next examined whether RSC-EXO also influences melanogenesis. To evaluate whether RSC-EXO affects melanogenesis, melanin production was assessed in α-MSH-stimulated B16F10 melanoma cells. Melanin content was quantified and normalized to cell viability to exclude cytotoxic effects.

RSC-EXO treatment resulted in reduction in melanin content at both tested concentrations, with decreases to 60% and 29% of control levels at low (3×10^8^ particles/mL) and high (1×10^9^ particles/mL) concentrations, respectively (Figure 2c). Notably, the inhibitory effect observed at the higher concentration was comparable to that of the positive control, arbutin (39%). In contrast, RSC-CM showed only minimal effects on melanin production (95% and 91% at low and high concentrations, respectively). Statistical significance was observed for the high-dose RSC-EXO treatment and the positive control, whereas other groups did not reach significance. These results suggest that RSC-EXO may modulate melanogenesis under the tested experimental conditions; however, this observation is based on a viability-normalized endpoint and should be interpreted with caution in the absence of further mechanistic validation, including analysis of melanogenesis-related pathways such as tyrosinase or MITF signaling.

### 2.5. Anti-inflammatory Activity of RSC-EXO

In addition to promoting tissue regeneration, effective skin therapeutics are expected to modulate inflammatory responses, which play a central role in skin damage and repair. Given the reported anti-inflammatory properties of *Rosa damascena*-derived materials, we next investigated whether RSC-EXO could regulate inflammatory signaling in immune cells.

Using an LPS-stimulated RAW 264.7 macrophage model, RSC-EXO treatment significantly suppressed the secretion of pro-inflammatory cytokines, including IL-1α, IL-6, and TNF-α, at both tested concentrations (Figure 3a–c). At 3 × 10^8^ particles/mL, RSC-EXO reduced IL-1α, IL-6, and TNF-α levels by approximately 72%, 75%, and 9%, respectively, while greater reductions were observed at 1 × 10^9^ particles/mL.

**Figure 3.**
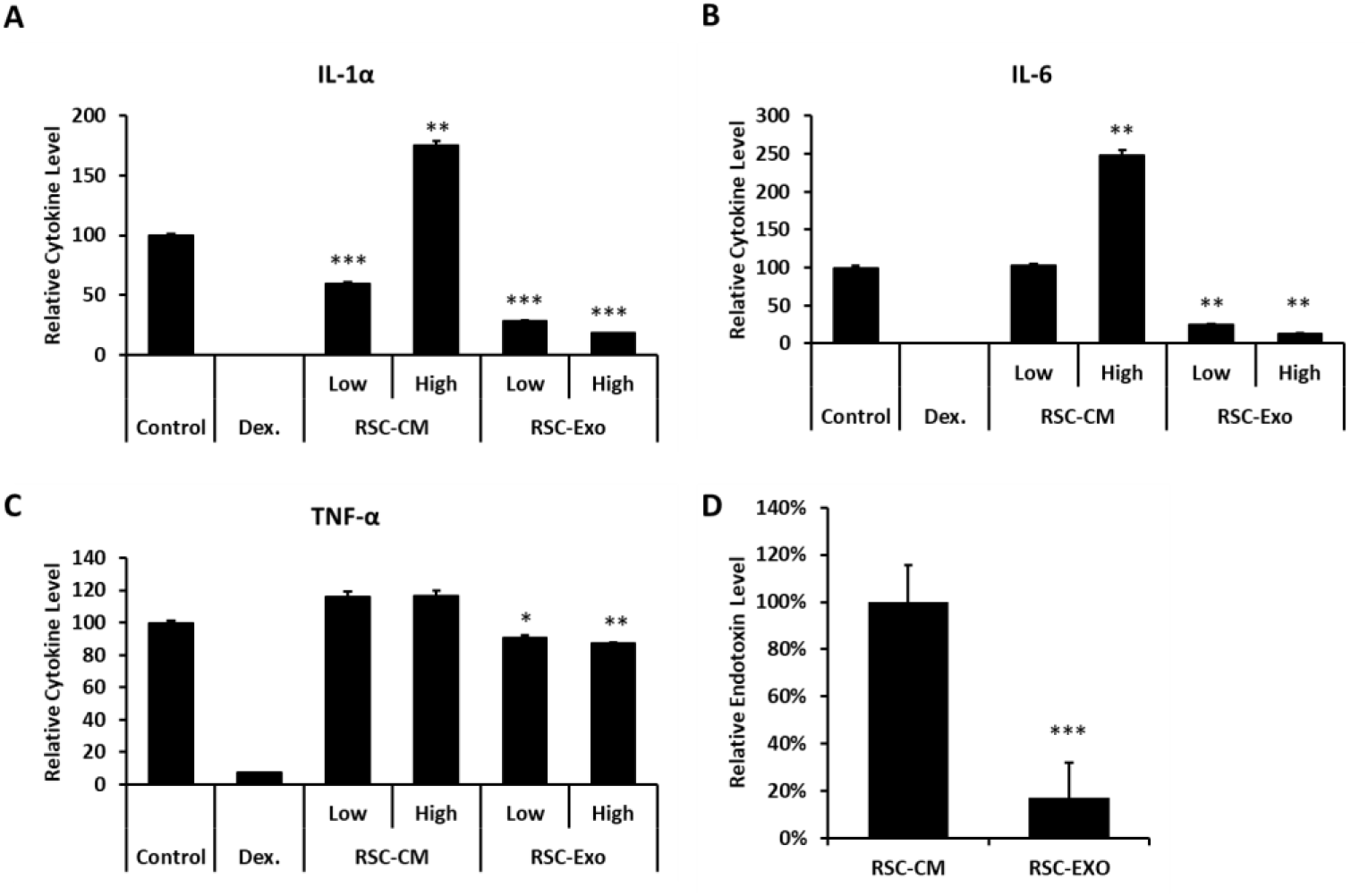
Anti-inflammatory effects of RSC-EXO. (A-C) RAW 264.7 murine macrophages were stimulated with lipopolysaccharide (LPS) to induce an inflammatory response and subsequently treated with RSC-EXO or RSC-CM. Pro-inflammatory cytokines IL-1α, IL-6, and TNF-α were quantified using the LEGENDplex^™^ Mouse Inflammation Panel (BioLegend, San Diego, CA, USA), a bead-based multiplex cytokine assay analyzed by flow cytometry. RSC-EXO was applied at high (1×10^9^ particles/mL) and low (3×10^8^ particles/mL) concentrations, whereas RSC-CM was treated at equivalent volumes. Dexamethasone (Dex) was used as a positive control, and PBS (vehicle control) was used as a negative control. (D) Relative endotoxin levels of RSC-CM and RSC-EXO measured by LAL assay. Endotoxin levels were normalized to RSC-CM (set as 100%). All experiments were performed in triplicate and repeated three times independently. Data are presented as mean ± SD (n = 3). Statistical significance was indicated as follows: \**p* < 0.05, \*\**p < 0.01, and* \*\*\**p < 0.001*.

In contrast, RSC-CM exhibited limited or inconsistent effects. Notably, RSC-CM increased IL-1α and IL-6 secretion at the tested concentrations, with statistically significant elevations observed at higher concentrations (except for IL-6 at the lower concentration), suggesting the presence of pro-inflammatory components in the crude conditioned medium, potentially including endotoxin-associated factors.

To further assess this possibility, endotoxin levels in RSC-CM and RSC-EXO were quantified using a LAL assay. When normalized to RSC-CM (set as 100%), RSC-EXO exhibited substantially lower endotoxin levels (17 ± 15%) compared to RSC-CM (100 ± 16%) (Figure 3d). These findings indicate that endotoxin-associated components may contribute to the pro-inflammatory effects observed in RSC-CM. However, despite the reduced endotoxin levels in RSC-EXO, treatment with RSC-EXO resulted in decreased pro-inflammatory cytokine levels compared to the vehicle control, suggesting that its anti-inflammatory effects cannot be solely attributed to differences in endotoxin content.

Collectively, these results suggest that RSC-EXO shows anti-inflammatory activity under the tested conditions, while crude conditioned medium may contain components that contribute to inflammatory responses. This interpretation should be considered with caution, as endotoxin-associated effects may partially influence the observed responses. This observation is consistent with the cytotoxicity profile described above and supports the notion that purification of vesicles enhances both biocompatibility and functional efficacy.

### 2.6. Proteomic Characterization of RSC-EXO

To gain molecular insight into the functional properties of RSC-EXO, proteomic profiling was performed. A total of 2,098 proteins were identified in the *Rosa damascena* callus and 206 proteins in RSC-EXO, among which 36 proteins were uniquely detected in the vesicle fraction (Figure 4a, Supplementary Data 1).

**Figure 4.**
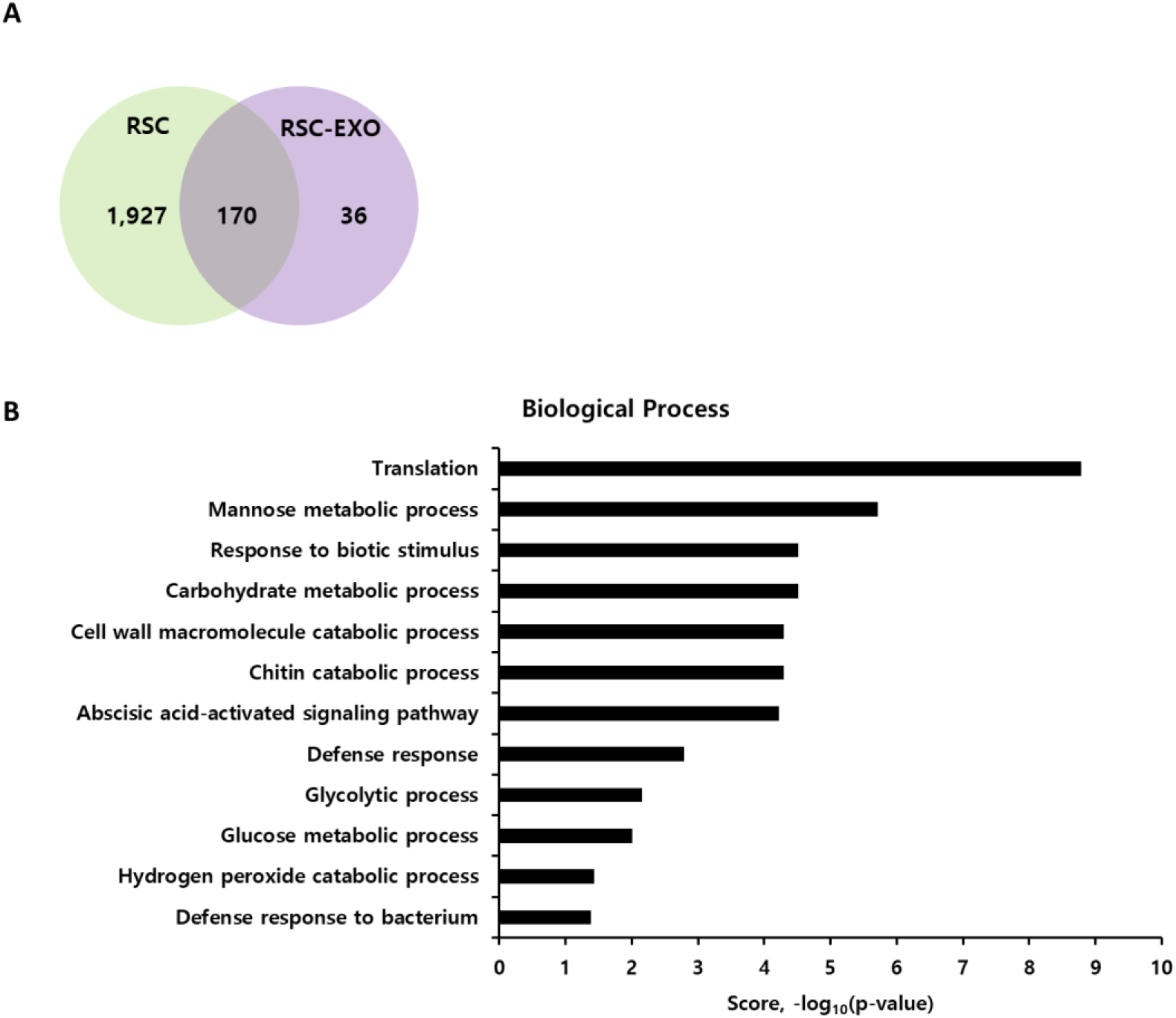
Proteomic analysis of RSC-EXO. (A) Venn diagram showing the number of proteins identified in *Rosa damascena* callus (RSC) and RSC-EXO, indicating proteins unique to each group or shared between them. (B) Gene Ontology (GO) enrichment analysis of proteins identified in RSC-EXO. Enriched biological processes included defense- and stimulus-responsive pathways, carbohydrate and cell wall macromolecule metabolism, and oxidative stress-related processes. GO terms for biological processes were analyzed using DAVID Bioinformatics Resources 6.8, and enrichment scores are expressed as –log_10_(p-value). Only terms with p < 0.05 (enrichment score ≥ 1.3) are shown.

Among the identified proteins, several relatively abundant proteins included peroxidases, Gnk2-like domaincontaining proteins, and START/Bet v I family proteins, which are associated with stress responses and defense-related processes in plants. In addition, multiple proteins related to cellular metabolism and energy production were also detected. These findings are consistent with the presence of a protein cargo associated with stress adaptation and core metabolic processes in RSC-EXO.

To further characterize the biological features of the identified proteins, Gene Ontology analysis was performed. The enriched biological processes could be broadly grouped into defense- and stimulus-responsive processes, carbohydrate and cell wall macromolecule metabolism, and core metabolic or oxidative stress-related pathways. In particular, terms such as response to biotic stimulus, defense response, defense response to bacterium, abscisic acid-activated signaling pathway, carbohydrate metabolic process, chitin catabolic process, cell wall macromolecule catabolic process, glycolytic process, glucose metabolic process, and hydrogen peroxide catabolic process were significantly represented in the RSC-EXO proteome (Figure 4b).

These findings indicate that the RSC-EXO proteome retains characteristic signatures of plant stress- and defense-associated biological processes. However, because the *Rosa damascena* proteome remains incompletely annotated, the specific functional roles of individual proteins identified in RSC-EXO will require further investigation.

### 2.7. Small RNA Profiling of RSC-EXO

Small RNA sequencing was performed to profile the small RNA population associated with RSC-EXO. A total of 75,897,704 raw reads were generated, with Q20 and Q30 values of 92.89% and 88.07%, respectively. After preprocessing, 46,402,609 reads were retained for downstream analysis (Supplementary Data 2). Read-length distribution analysis revealed a detectable fraction of small RNAs within the canonical miRNA size range (18–25 nt), accounting for approximately 8.0% of the processed reads (Figure 5). In contrast, a substantial proportion of reads was observed outside this range, with a dominant peak at ~48 nt, indicating the presence of longer RNA fragments. These results suggest that RSC-EXO contains a heterogeneous population of small RNAs, including a limited fraction of miRNA-sized reads together with longer RNA fragments.

**Figure 5.**
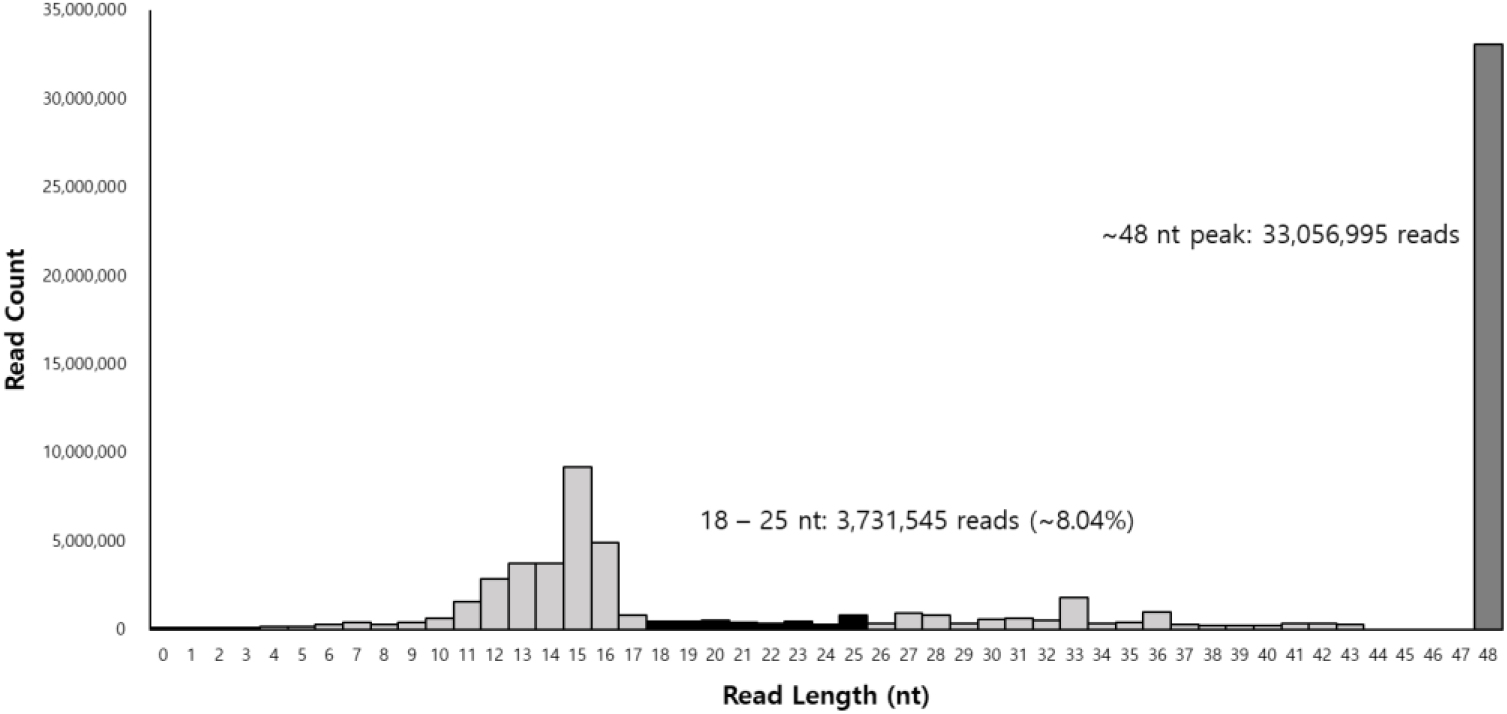
Read length distribution of small RNA sequences in RSC-EXO. The distribution of read lengths obtained from small RNA sequencing is shown. A fraction of reads was observed within the canonical miRNA size range (18–25 nt), representing approximately 8.0% of processed reads. A dominant peak was detected at ~48 nt, indicating the presence of longer RNA fragments. These results suggest that RSC-EXO contains a heterogeneous population of small RNAs, including a limited fraction of miRNA-sized reads together with longer RNA fragments.

To assess genome alignment, processed reads were mapped to the *Rosa damascena* reference genome. A subset of reads (5.16%) was successfully mapped to the reference genome, whereas a large proportion remained unmapped. Consistently, RNA composition analysis indicated that most reads were classified as unknown, with smaller fractions assigned to genomic regions and predicted miRNA-like candidate sequences. This pattern is likely attributable to the limited annotation of Rosa small RNAs and the presence of diverse non-canonical or fragmented RNA species in plant-derived extracellular vesicles.

Genome-based prediction identified 39 precursor and 36 mature candidate small RNA sequences with predicted miRNA-like features (Supplementary Data 3). Among these, a subset of candidates meeting predefined confidence criteria (miRDeep2 score ≥ 1, significant randfold p-value, and mature read count ≥ 100) showed relatively high read abundance and predicted secondary structures compatible with miRNA-like features, including detectable star strand reads (Supplementary Data 4). Representative predicted structures are shown in Supplementary Data 5.

Overall, these results indicate that RSC-EXO contains a heterogeneous population of small RNAs, including a limited fraction of miRNA-sized reads, while a considerable proportion of sequences remains unannotated. Therefore, the current small RNA profiling data should be interpreted as exploratory rather than definitive characterization of functional miRNA cargo.

## 3. Discussion

In this study, we report the isolation and characterization of exosome-like vesicles derived from the culture medium of *Rosa damascena* callus (RSC-EXO). These vesicles exhibited key physicochemical and molecular features consistent with extracellular vesicles (EVs), including nanoscale size distribution, spherical morphology, enrichment of plant EV-associated markers (PEN1 and TET8), and efficient uptake by human dermal fibroblasts (HDFs). While NTA, cryo-TEM, and marker analysis support the presence of vesicle-like structures, the current characterization does not fully meet the stringent criteria for defining a highly purified exosome population. Compared with previous reports that primarily focused on crude plant extracts or conditioned media, this study provides a characterization of vesicle-enriched fractions derived from plant callus systems, thereby contributing to the growing body of research on plant-derived extracellular vesicles (PDEVs) as biologically active entities [9–11, 16, 17]. Importantly, this study provides an integrated molecular and functional characterization of callus-derived plant extracellular vesicles from Rosa damascena, a system that has remained insufficiently explored. The vesicle population characterized in this study is referred to as exosome-like vesicles, as the current data do not fully meet the stringent criteria required for defining highly purified exosomes.

Consistent with previous studies, plant-derived EVs have been reported to mediate intercellular communication and to interact with mammalian systems through cellular uptake mechanisms such as endocytosis or membrane fusion [14, 40–42]. Their structural stability under physiological and environmental conditions further supports their potential applicability in topical or systemic delivery contexts [11–13].

Functionally, RSC-EXO showed a consistent ability to enhance collagen synthesis and to promote wound closure-related cellular responses in HDFs. While the increase in wound closure did not reach statistical significance, a reproducible trend toward enhanced wound closure was observed, suggesting a potential contribution to dermal regeneration processes. However, because the scratch assay used in this study does not fully distinguish between cell migration and proliferation, both processes may have contributed to the observed wound closure response. Importantly, these functional effects were accompanied by a markedly improved safety profile compared to the crude conditioned medium (RSC-CM). When normalized by particle number, RSC-EXO exhibited substantially lower cytotoxicity, indicating superior biocompatibility.

In addition to its lower cytotoxicity, RSC-EXO showed anti-inflammatory activity, whereas RSC-CM showed proinflammatory tendencies at the tested concentrations, particularly in IL-1α and IL-6 secretion. Together, these findings suggest that the applied purification strategy, involving multi-step separation, may improve biocompatibility while reducing potentially detrimental components present in the bulk conditioned medium. It should be noted that endotoxin-associated components present in plant-derived conditioned media may contribute to macrophage activation, and therefore the observed effects should be interpreted with caution. However, the reduction in cytokine levels observed with RSC-EXO compared to the vehicle control suggests that its anti-inflammatory effects cannot be solely attributed to endotoxin differences.

Beyond dermal regeneration, RSC-EXO also exhibited a modulatory effect on melanogenesis, as evidenced by reduced melanin production in α-MSH-stimulated melanoma cells. Although this observation is based on viability-normalized endpoint measurements and requires further mechanistic validation, it suggests that RSC-EXO may influence multiple aspects of skin physiology, including pigmentation.

Proteomic analysis revealed that RSC-EXO contains a distinct subset of proteins compared with the parental callus, indicating selective cargo loading during vesicle biogenesis. The identified proteins were broadly associated with stress responses, defense mechanisms, carbohydrate metabolism, and oxidative processes, suggesting that RSC-EXO retains characteristic molecular signatures of plant stress-adaptive systems. Notably, the enrichment of Gene Ontology terms such as response to biotic stimulus, defense response, and oxidative stress-related processes is consistent with previous reports demonstrating that plant-derived extracellular vesicles are enriched in proteins involved in defense signaling, stress adaptation, and extracellular communication [18–20]. However, these annotations originate from plant-specific biological contexts and should not be directly interpreted as equivalent to mammalian inflammatory or regenerative pathways. It should also be noted that protein identification was performed using the *Rosa chinensis* reference database due to the limited annotation available for *Rosa damascena*. This may influence protein assignment and functional interpretation, and therefore the proteomic results should be interpreted with caution.

Small RNA profiling revealed a heterogeneous RNA population within RSC-EXO, including only a limited fraction of sequences within the canonical miRNA size range. In contrast, a dominant peak was observed at approximately 48 nt, and only a small proportion of reads mapped to the *Rosa damascena* reference genome. A substantial proportion of reads remained unannotated, which may reflect current limitations in *Rosa* small RNA databases as well as the presence of diverse non-canonical or fragmented RNA species, including possible tRNA/rRNA-derived fragments. Previous studies have highlighted challenges in plant miRNA annotation, particularly in non-model species, and have also reported the heterogeneous nature of small RNA populations associated with plant-derived extracellular vesicles [18, 21, 26, 42, 43]. Therefore, the present findings should be interpreted as exploratory profiling of heterogeneous small RNA populations rather than definitive evidence of functional miRNA cargo.

Collectively, these findings suggest that RSC-EXO exhibits multiple biological activities under the tested in vitro conditions, including enhancement of collagen synthesis, modulation of wound-related cellular responses, regulation of melanogenesis, and suppression of inflammatory cytokines. Importantly, these effects were consistently observed in the purified vesicle fraction but not in the crude conditioned medium, underscoring the significance of vesicle isolation in defining functional bioactivity. This integrated functional and molecular characterization further supports the potential of plant callus-derived extracellular vesicles as a reproducible and scalable bioactive platform.

Despite these promising findings, several limitations should be acknowledged. First, the observed biological effects are based on in vitro assays and require validation in in vivo systems. Second, the molecular mechanisms underlying the observed effects remain incompletely defined, and no loss-of-function or gain-of-function experiments were performed to establish causal relationships between specific protein or RNA cargo and the observed biological responses. Third, the limited annotation of *Rosa* genomic and proteomic databases constrains the depth of molecular interpretation. In addition, the functional assays were performed using a limited concentration range, and therefore definitive dose-response relationships will require further investigation.

In conclusion, this study suggests that RSC-EXO represents a biologically active plant-derived extracellular vesicle population with regenerative and anti-inflammatory activity observed in human skin-related cellular models. The findings highlight the importance of vesicle purification in enhancing both biocompatibility and functional efficacy and suggest that plant callus-derived EVs may serve as a promising platform for future cosmeceutical and dermatological applications. Further studies focusing on in vivo validation, molecular mechanism elucidation, and optimization of delivery systems will be essential to fully realize the translational potential of RSC-EXO.

## 4. Materials and Methods

### 4.1. Callus Culture and Collection of Conditioned Medium

Callus tissues were induced from petals of Rosa damascena. Sterilized flower buds were treated with 70% ethanol and sodium hypochlorite, rinsed with sterile water, and the excised petals were placed on Murashige and Skoog (MS) medium supplemented with 2,4-dichlorophenoxyacetic acid (2,4-D) and kinetin to induce callus formation. Callus tissues were maintained in the dark at 25 °C and subcultured every two weeks on the same medium. To minimize culture-dependent variability, callus tissues were maintained under the same induction medium, subculture interval, suspension culture conditions, and harvest timing throughout the study. For suspension culture, proliferating callus tissues were transferred to liquid MS medium containing 3% sucrose and 1 mg/L 2,4-D (pH 5.8) and cultured at 25 °C under dark conditions with gentle agitation. After 20 days, the conditioned medium (RSC-CM) was collected and sequentially centrifuged at 300 × g for 10 min, 2,000 × g for 20 min, and 10,000 × g for 30 min to remove cells and debris, followed by filtration through a 0.22 μm membrane.

### 4.2. Purification and Nanoparticle Tracking Analysis of RSC-EXO

RSC-EXO were initially isolated from rose callus-conditioned medium (RSC-CM) using a tangential flow filtration (TFF)-based approach, as previously described for mammalian stem cell-derived exosomes [44]. Briefly, filtered RSCCM was concentrated and diafiltrated with phosphate-buffered saline (PBS). To enhance vesicle purity and reduce nonvesicular contaminants, the TFF-isolated fraction was further processed using a sequential multi-step chromatographic purification implemented on an FPLC platform, combining multimodal chromatography and ion-exchange membrane adsorption steps (Capto Core multimodal resin, Cytiva, USA, and Sartobind Q anion-exchange membrane, Sartorius, Germany). This strategy integrates multiple physicochemical separation modalities, including size-based and charge-based separation mechanisms, to refine vesicle-enriched fractions. The purification workflow was designed to minimize co-isolated soluble proteins and non-vesicular particles while preserving vesicle integrity. Purified RSC-EXO were stored at −80 °C until analysis.

Particle size distribution and concentration were determined using a NanoSight NS300 system (Malvern Panalytical, Amesbury, UK). Three independent measurements were performed for each sample, and averaged values were used to determine mean diameter, mode diameter, and particle concentration.

### 4.3. Cryogenic Transmission Electron Microscopy

The morphology of RSC-EXO was examined by cryogenic transmission electron microscopy (Cryo-TEM). Purified vesicles were applied to holey carbon grids, vitrified in liquid ethane, and imaged under cryogenic conditions using a transmission electron microscope (JEOL JEM-2100Plus, JEOL Ltd., Tokyo, Japan) operated at 200 kV.

### 4.4. Western Blot Analysis of RSC-EXO

Equal amounts of total protein (10 μg) from rose callus lysate, RSC-CM, and RSC-EXO were subjected to Western blot analysis. Proteins were extracted in RIPA buffer containing protease inhibitors, separated by SDS-PAGE, and transferred to PVDF membranes.

After blocking, membranes were incubated overnight at 4 °C with primary antibodies against PEN1 (1:500; Cusabio, CSB-PA875527XA01DOA, Wuhan, China), TET8 (1:1000; PhytoAB, PHY1490A, San Jose, CA, USA), and GAPDH (1:1000; MyBioSource, MBS9373474, Rosemont, IL, USA), followed by HRP-conjugated secondary antibodies (Jackson ImmunoResearch, West Grove, PA, USA). Protein bands were visualized using an enhanced chemiluminescence (ECL) reagents (DoGenBio, Seoul, Republic of Korea) and imaged using an Amersham Imager 680 system (GE Healthcare, Chicago, IL, USA). Uncropped Western blot images are provided in Supplementary Data S6.

### 4.5. Cellular Uptake of RSC-EXO

Purified RSC-EXO were labeled with PKH67 Green Fluorescent Cell Linker (Sigma-Aldrich, St. Louis, MO, USA) according to the manufacturer’s instructions. Excess unbound dye was removed using MiniTrap G-25 desalting columns (Sartorius, Göttingen, Germany).

As a control for dye-related artifacts, PBS was subjected to the same PKH67 labeling and purification procedure in the absence of RSC-EXO (dye-only control).

Human dermal fibroblasts (HDFs; ATCC, Manassas, VA, USA) were incubated with PKH67-labeled RSC-EXO or dye-only control for 24 or 48 h at 37 °C. Cells were then fixed with 10% neutral-buffered formalin, stained with Hoechst 33258 for nuclei and CellMask Deep Red for cytoplasm, and analyzed by fluorescence microscopy.

### 4.6. Cytotoxicity Assay

The cytotoxicity of RSC-EXO and RSC-CM was evaluated in HDFs using an MTT assay. Cells were seeded in 96-well plates at a density of 1 × 10^4^ cells/well and cultured for 24 h. Cells were then treated with the indicated concentrations of RSC-EXO or equivalent volumes of RSC-CM for 24 h.

After treatment, cells were incubated with MTT reagent, and the resulting formazan crystals were dissolved in dimethyl sulfoxide (DMSO). Absorbance was measured at 570 nm, and cell viability was expressed as a percentage relative to the untreated control.

### 4.7. Scratch Wound Assay

HDF migration was assessed using a scratch-wound assay with the Incucyte® Live-Cell Imaging System (Sartorius, Göttingen, Germany). HDFs were seeded in 96-well plates and cultured to confluence. A uniform linear scratch was introduced using the WoundMaker^™^, debris was removed by washing with Dulbecco’s phosphate-buffered saline (DPBS), and cells were treated with RSC-CM or RSC-EXO under serum-free conditions. A medium containing 5% FBS was used as a positive control. Wound closure was monitored by time-lapse imaging for 24 h.

### 4.8. Collagen Assay

Procollagen type I C-peptide (PIP) levels were quantified using a commercial ELISA kit (Takara, Shiga, Japan) according to the manufacturer’s instructions. HDFs were seeded in 24-well plates at 2.5 × 10^4^ cells/well, incubated for 24 h, and treated with RSC-EXO or RSC-CM at the indicated concentrations.

After treatment, culture supernatants were collected and transferred to an ELISA plate for PIP quantification according to the manufacturer’s protocol. The ELISA reaction was stopped with stop solution, and absorbance was measured at 450 nm. After removal of the culture supernatants, the remaining cells in the original culture plate were subjected to an MTT assay, and PIP levels were normalized to the corresponding cell viability readout.

### 4.9. Melanin Content Assay

B16F10 murine melanoma cells (ATCC) were cultured in phenol red-free DMEM supplemented with 10% FBS, penicillin (50 U/mL), streptomycin (50 μg/mL), 2 mM L-glutamine, and 1 mM sodium pyruvate. Cells were seeded in 48-well plates at a density of 8.0 × 10^3^ cells/well and incubated for 24 h at 37 °C under 5% CO_2_.

The medium was then replaced with fresh medium containing RSC-EXO or RSC-CM, and cells were further incubated for 48 h. To induce melanogenesis, α-melanocyte-stimulating hormone (α-MSH; Sigma-Aldrich) was added to all groups at a final concentration of 100 nM. Arbutin (1 mM; Sigma-Aldrich) was used as a positive control.

After treatment, extracellular melanin in the culture supernatant and intracellular melanin were collected separately. Cell viability was assessed using a CCK-8 assay (Dojindo, Kumamoto, Japan) at 450 nm. Cells were then washed with DPBS and lysed in 1 N ammonium hydroxide containing 10% DMSO at 85 °C for 30 min to extract intracellular melanin.

Melanin content was quantified at 405 nm using a synthetic melanin standard curve, and the measured melanin values were normalized to cell viability.

### 4.10. Measurement of Inflammatory Cytokines

RAW 264.7 macrophages (ATCC, Manassas, VA, USA) were seeded at 3 × 10^4^ cells/well in 48-well plates and treated with RSC-CM or RSC-EXO in the presence of lipopolysaccharide (LPS, 10 ng/mL).

After incubation, culture supernatants were collected, and the concentrations of IL-1α, IL-6, and TNF-α were quantified using the LEGENDplex^™^ Mouse Inflammation Panel (BioLegend, San Diego, CA, USA) according to the manufacturer’s instructions. Samples were analyzed using a NovoCyte 2000 flow cytometer (Agilent Technologies, Santa Clara, CA, USA), and data were processed using LEGENDplex^™^ Data Analysis Software.

### 4.11. Proteomic Analysis of RSC-EXO

Total proteins were extracted from purified RSC-EXO, quantified using a bicinchoninic acid (BCA) assay, separated by SDS-PAGE, and subjected to in-gel tryptic digestion. Peptides were analyzed by liquid chromatography–tandem mass spectrometry (LC–MS/MS) using a Q Exactive Plus mass spectrometer (Thermo Fisher Scientific, Waltham, MA, USA) in data-dependent acquisition mode.

Protein identification was performed using MASCOT version 2.7 (Matrix Science, London, UK) against the Rosa chinensis protein database, due to the limited availability of a fully annotated Rosa damascena proteome. A false discovery rate (FDR) threshold of 1% was applied.

Gene Ontology (GO) enrichment analysis for Biological Process terms was conducted using DAVID Bioinformatics Resources 6.8. Only GO terms with p < 0.05 were considered significant.

### 4.12. Small RNA Profiling of RSC-EXO

Total RNA was extracted from purified RSC-EXO using the miRNeasy Mini Kit (Qiagen, Hilden, Germany). RNA quantity and quality were assessed spectrophotometrically.

Small RNA libraries were prepared using the SMARTer smRNA-Seq Kit (Takara, Shiga, Japan) and sequenced on an Illumina platform (single-end, 51 bp). Raw reads were adapter-trimmed and quality-filtered, and processed reads were analyzed for length distribution.

Processed reads were mapped to the Rosa damascena reference genome (NCBI assembly ASM166254v1), and candidate small RNA sequences with miRNA-like features were predicted using a miRDeep2-based pipeline. Candidate features including read abundance, hairpin structure, randfold significance, and star strand support were evaluated.

### 4.13. Statistical Analysis

All data are presented as mean ± standard deviation (SD) from at least three independent experiments. Statistical analyses were performed using GraphPad Prism version 8.4.3 (GraphPad Software, San Diego, CA, USA). Differences between groups were evaluated by one-way analysis of variance (ANOVA) followed by Sidak’s post hoc test. A p-value < 0.05 was considered statistically significant.

## Supplementary Materials

Supplementary Data 1: RSC-EXO Protein Profile; Supplementary Data 2: RSC-EXO small RNA sequencing and preprocessing summary; Supplementary Data 3. Predicted small RNA candidates with miRNA-like features identified in RSC-EXO; Supplementary Data 4. High-confidence candidate small RNA sequences identified in RSC-EXO; Supplementary Data 5. Representative predicted structures of candidate small RNAs in RSC-EXO; Supplementary Data 6. Uncropped Western blot images.

## Author Contributions

Conceptualization, B.S.C., and D.H.H.; methodology, E.L.; validation, H.J.L., S.Y.M., E.S., and B.S.; formal analysis, E.L.; investigation, H.J.L., S.Y.M., E.S., B.S., J.J.L., J.Y.H., S.K.P; data curation, E.L., K.L.; writing—original draft preparation, E.L. and D.H.H.; writing—review and editing, C.P., and M.S.C.; supervision, B.S.C., D.H.H., and M.S.C.; project administration, B.S.C., D.H.H., and M.S.C.

All authors have read and agreed to the published version of the manuscript.

## Institutional Review Board Statement

Not applicable

## Informed Consent Statement

Not applicable

## Data Availability Statement

The raw data supporting the conclusions of this article will be made available by the authors on request.

## Acknowledgments

During the preparation of this manuscript, the authors used ChatGPT (OpenAI, GPT-5, 2025) for the purpose of English language editing. The authors have carefully reviewed and revised the output and take full responsibility for the content of this publication.

