## Supplementary material for "Characterization of *Rosa damascena* Callus-derived Exosome-like Vesicles and Their Multifunctional Activities in Skin-related Cellular Models": Supplementary Data 5. Representative predicted structures of candidate small RNAs in RSC-EXO.pdf

Two representative candidate small RNA sequences (LYNE01003137_18835 and LYNE01061673_42069) identified through exploratory computational analysis are shown. These candidates exhibited relatively high read abundance and predicted secondary structures compatible with miRNA-like features; however, their biological function and classification remain unvalidated.

| | | |
|---|---|---|
| Provisional ID | : | LYNE01003137_18835 |
| Score total | : | 619.1 |
| Score for star read(s) | : | 3.9 |
| Score for read counts | : | 612.4 |
| Score for mfe | : | 1.1 |
| Score for randfold | : | 1.6 |
| Score for cons. seed | : | |
| Total read count | : | 1213 |
| Mature read count | : | 1200 |
| Loop read count | : | 0 |
| Star read count | : | 13 |

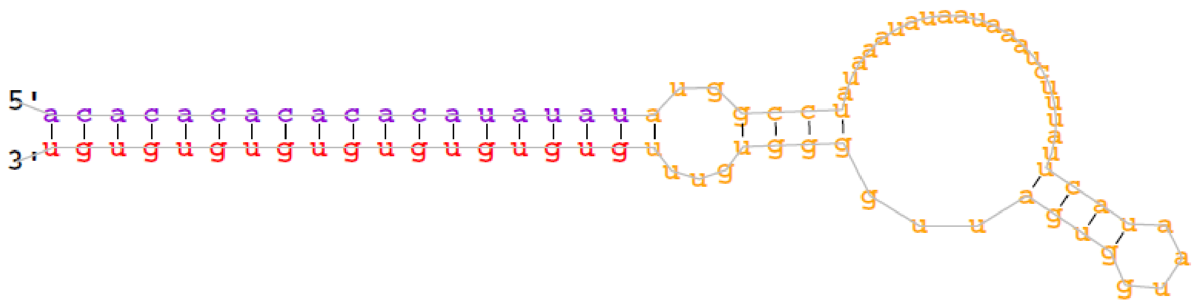


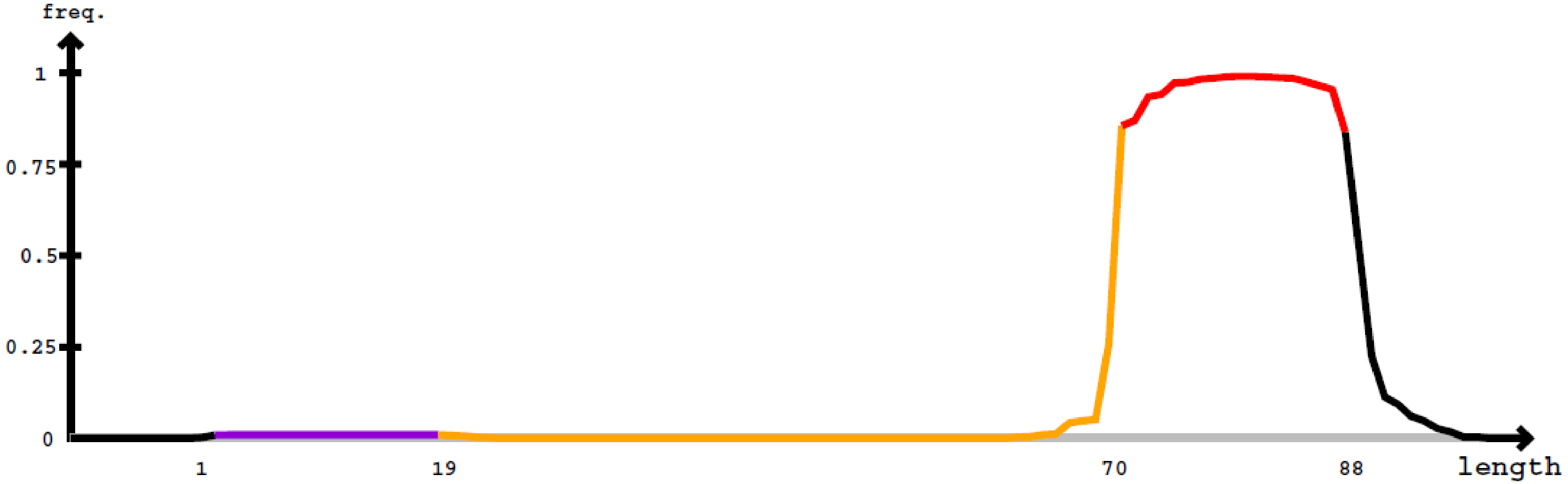


Star Mature

5'- uuucauuuaacacacacacacauauauauggccuauaaauauaauaaaucuuuauucauaauggugauugggguguuugugugugugugugugugu auguguguauuaa -3' obs

uuucauuuaacacacacacacauauauauggccuauaaauauaauaaaucuuuauucauaauggugauugggguguuugugugugugugugugugu auguguguauuaa exp

| | | |
|---|---|---|
| Provisional ID | : | LYNE01061673_42069 |
| Score total | : | 580.8 |
| Score for star read(s) | : | 3.9 |
| Score for read counts | : | 573.7 |
| Score for mfe | : | 1.6 |
| Score for randfold | : | 1.6 |
| Score for cons. seed | : | |
| Total read count | : | 1137 |
| Mature read count | : | 1123 |
| Loop read count | : | 0 |
| Star read count | : | 14 |

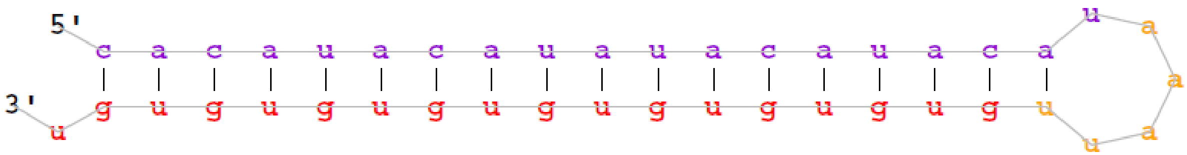


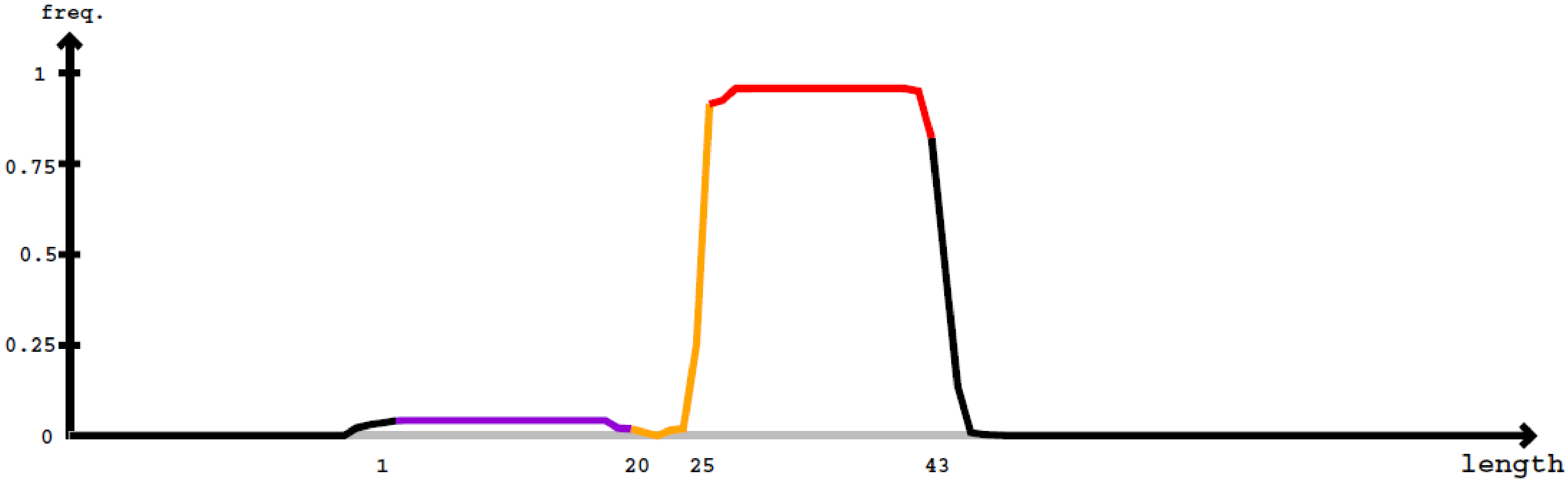


Star Mature

5'- aaaggugaauaacugacuagcuacacauacauauacauacauaaauugugugugugugugugugugu auuacauauguauauugacaugcaguuuuuucauuuuauuuugau -3' obs

aaaggugaauaacugacuagcuacacauacauauacauacauaaauugugugugugugugugugugu auuacauauguauauugacaugcaguuuuuucauuuuauuuugau exp
