## Supplementary material for "Characterization of *Rosa damascena* Callus-derived Exosome-like Vesicles and Their Multifunctional Activities in Skin-related Cellular Models": Supplementary Data 5. Representative predicted structures of candidate small RNAs in RSC-EXO.pptx

### Slide 1
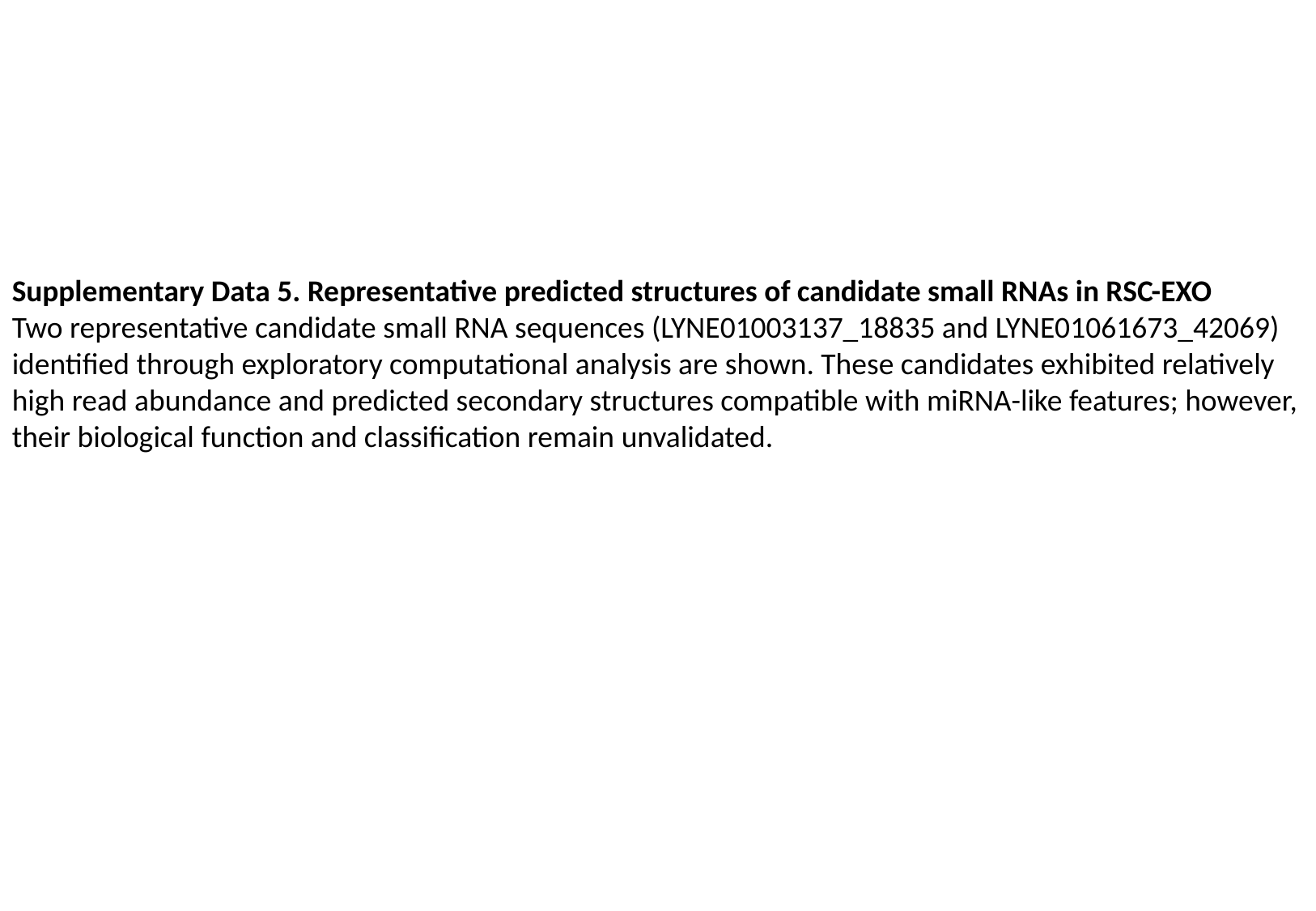

Supplementary Data 5. Representative predicted structures of candidate small RNAs in RSC-EXO
Two representative candidate small RNA sequences (LYNE01003137_18835 and LYNE01061673_42069) identified through exploratory computational analysis are shown. These candidates exhibited relatively high read abundance and predicted secondary structures compatible with miRNA-like features; however, their biological function and classification remain unvalidated.

### Slide 2
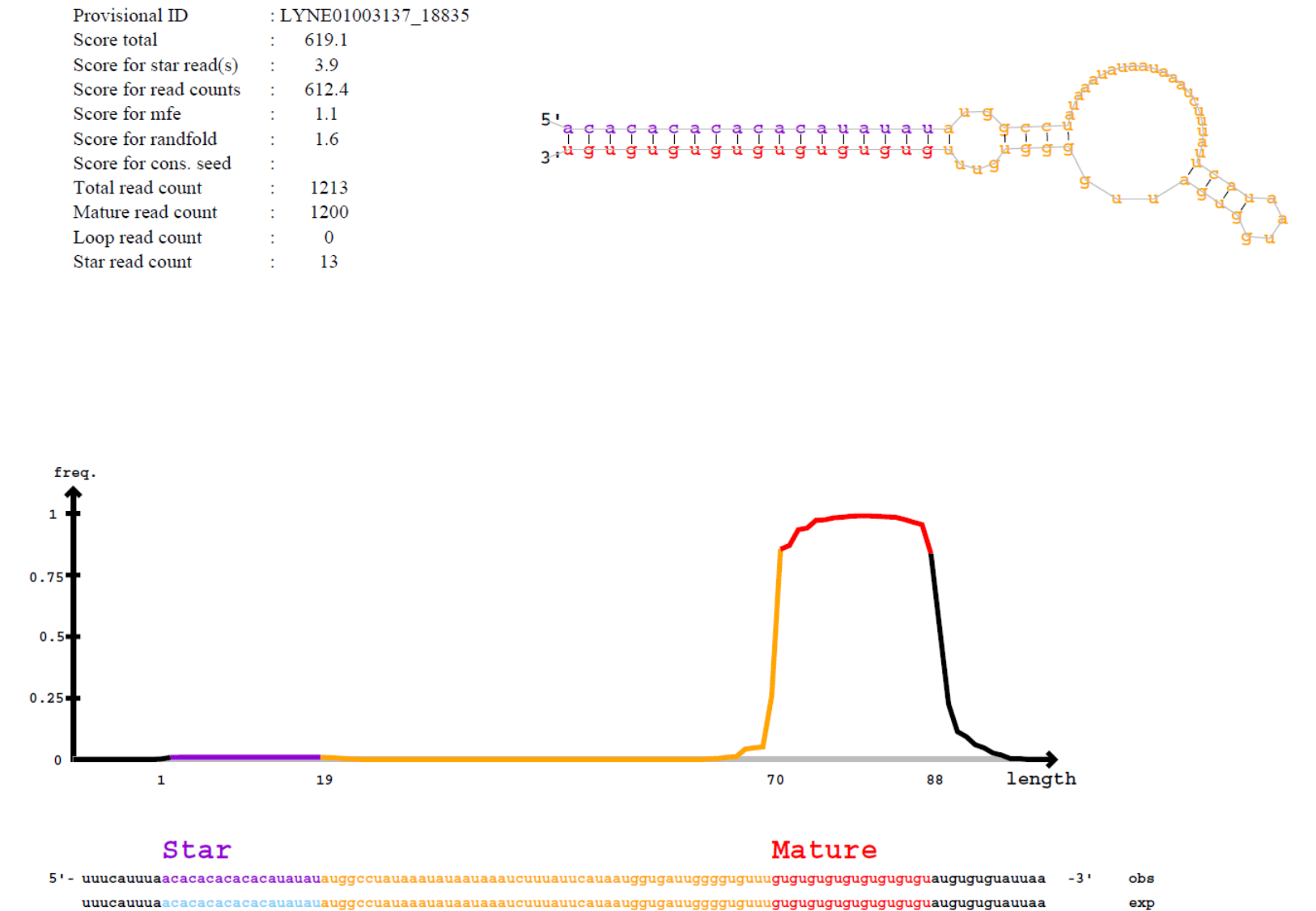

### Slide 3
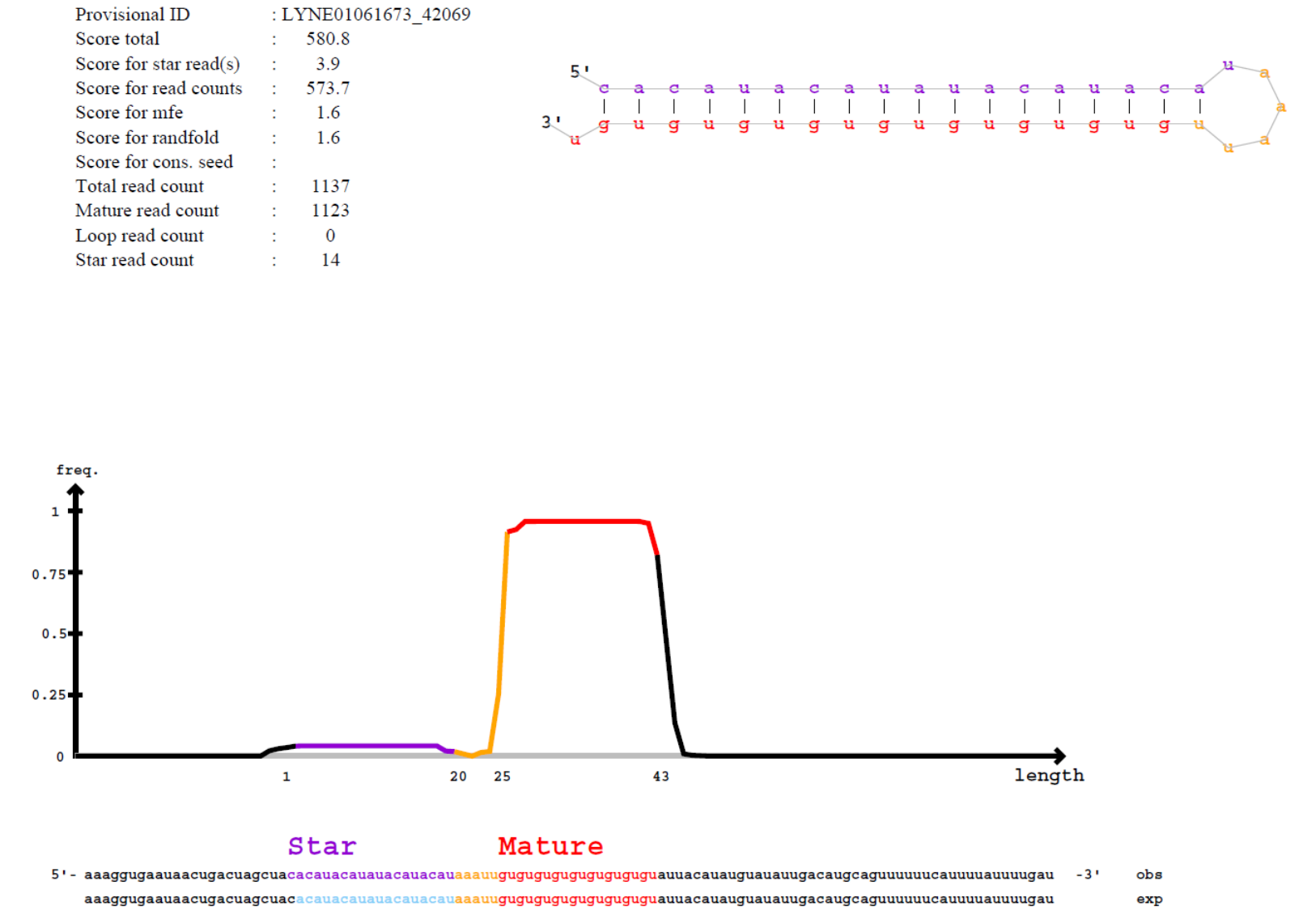
