## Supplementary material for "Characterization of *Rosa damascena* Callus-derived Exosome-like Vesicles and Their Multifunctional Activities in Skin-related Cellular Models": Supplementary Data 6. Uncropped Western blot images.pdf

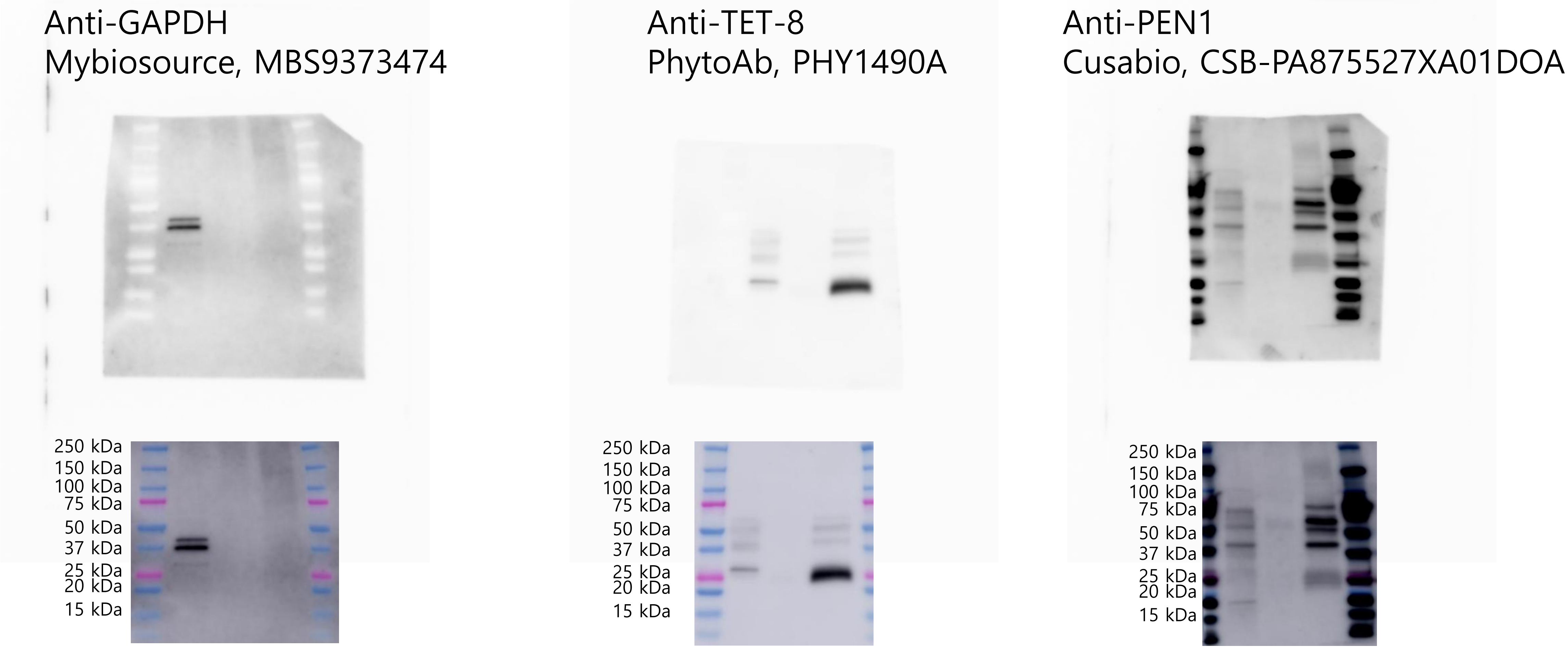


**Supplementary Data 6. Uncropped Western blot images.**
Uncropped Western blot images corresponding to Figure 1C. The full-length blots are shown with molecular weight markers and lane annotations. These images represent the original, unprocessed data prior to cropping for presentation in the main figures.
