## Supplementary material for "Characterization of *Rosa damascena* Callus-derived Exosome-like Vesicles and Their Multifunctional Activities in Skin-related Cellular Models": Supplementary Data 6. Uncropped Western blot images.pptx

### Slide 1
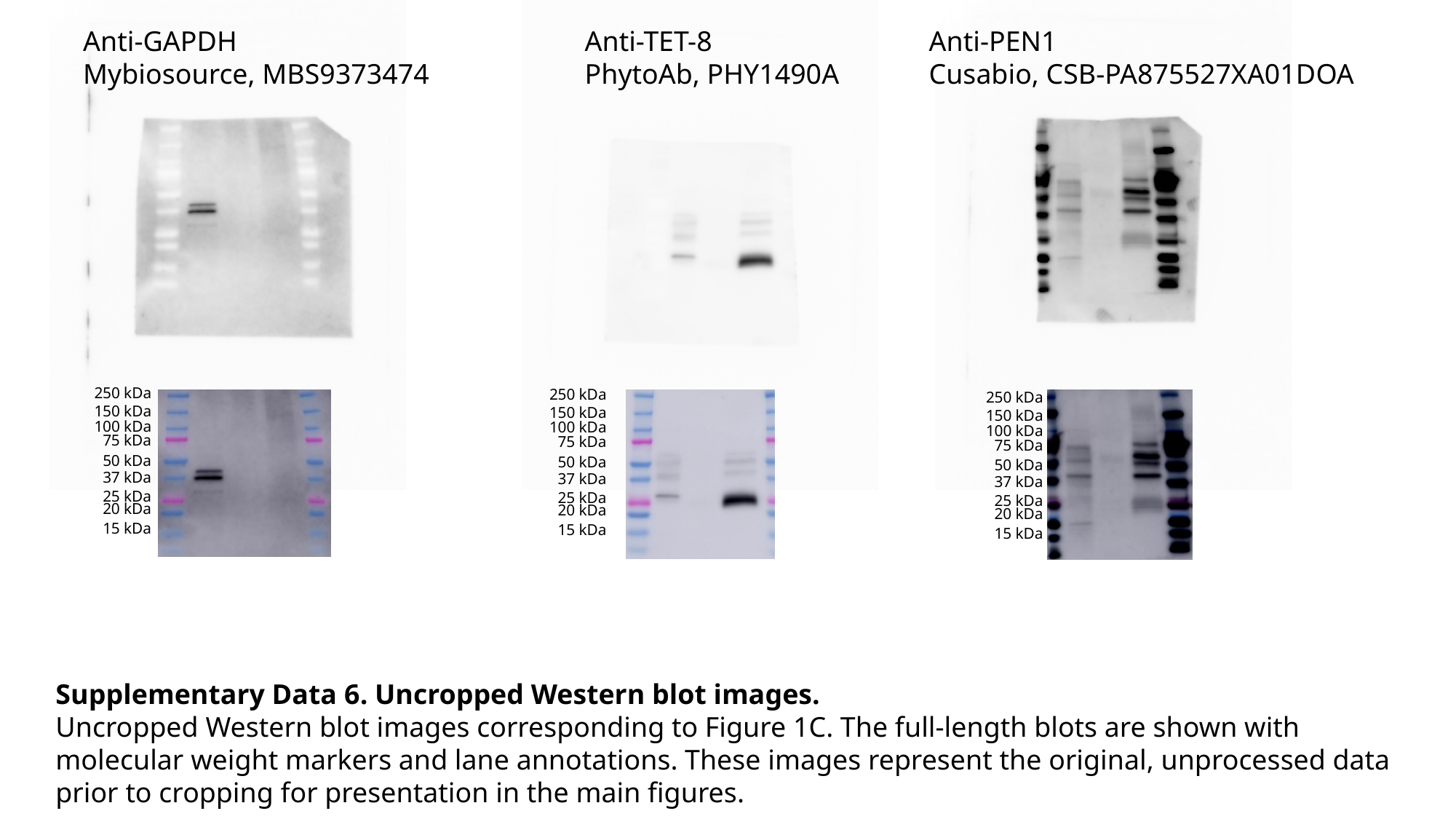

Anti-TET-8
PhytoAb, PHY1490A
Anti-PEN1
Cusabio, CSB-PA875527XA01DOA
Anti-GAPDH
Mybiosource, MBS9373474
250 kDa
250 kDa
250 kDa
150 kDa
150 kDa
150 kDa
100 kDa
100 kDa
100 kDa
75 kDa
75 kDa
75 kDa
50 kDa
50 kDa
50 kDa
37 kDa
37 kDa
37 kDa
25 kDa
25 kDa
25 kDa
20 kDa
20 kDa
20 kDa
15 kDa
15 kDa
15 kDa
Supplementary Data 6. Uncropped Western blot images.
Uncropped Western blot images corresponding to Figure 1C. The full-length blots are shown with molecular weight markers and lane annotations. These images represent the original, unprocessed data prior to cropping for presentation in the main figures.
